# The interaction between non-fusogenic Sec22b-Stx complexes and Extended-Synaptotagmins promotes neurite growth and ramification

**DOI:** 10.1101/2019.12.24.886648

**Authors:** Alessandra Gallo, Lydia Danglot, Francesca Giordano, Thomas Binz, Christian Vannier, Thierry Galli

## Abstract

Axons and dendrites are long and often ramified neurites that need particularly intense plasma membrane (PM) expansion during the development of the nervous system. Neurite growth depends on non-fusogenic Sec22b–Stx1 SNARE complexes at endoplasmic reticulum (ER)-PM contacts. Here we show that Sec22b interacts with the endoplasmic reticulum lipid transfer proteins Extended-Synaptotagmins (E-Syts) and this interaction depends on the Longin domain of Sec22b. Overexpression of E-Syts stabilizes Sec22b-Stx1 association, whereas silencing of E-Syts has the opposite effect. Overexpression of wild-type E-Syt2, but not mutants unable to transfer lipids or attach to the ER, increase the formation of axonal filopodia and ramification of neurites in developing neurons. This effect is inhibited by a clostridial neurotoxin cleaving Stx1, expression of Sec22b Longin domain and a Sec22b mutant with extended linker between SNARE and transmembrane domains. We conclude that Sec22b-Stx1 ER-PM contact sites contribute to PM expansion by interacting with lipid transfer proteins such as E-Syts.

## Introduction

Tissue and organism growth is supported by the growth of each cell after each cell division. Plasma membrane (PM) and intracellular membranes growth support cell growth. Cell growth is particularly intense in highly polarized cells like neurons. During their development, neurons elaborate processes extending hundreds of microns to meters from the cell body, requiring an increase in their PM surface by 20% per day (Pfenninger, 2009). Hence, compared to other cell types, developing neurons have to face a formidable challenge of adding new membrane to appropriate locations in a manner that requires both high processivity and fine regulation.

Membrane expansion during neuronal development has been thought to be mediated by soluble N-ethylmalemide-sensitive attachment protein receptor (SNARE)-dependent fusion of secretory vesicles with the PM (Wojnacki & Galli, 2016). SNAREs are transmembrane proteins mediating membrane fusion in all the trafficking steps of the secretory pathway. In order for fusion to occur, a *trans*-SNARE complex, composed of three Q-SNAREs (on the acceptor compartment) and one R-SNARE (on the opposing membrane), assemble to bring the opposite lipid bilayers in close proximity and trigger their fusion (Jahn & Scheller, 2006; Südhof & Rothman, 2009). In mammals, the R-SNAREs Syb2/VAMP2, VAMP4 and TI-VAMP/VAMP7 have been implicated in neurite extension (Alberts *et al*, 2003; Martinez-Arca *et al*, 2001; Grassi *et al*, 2015; Schulte *et al*, 2010). However, single knockouts (KO) mice for VAMP7 (Danglot *et al*, 2012) or VAMP2 (Schoch *et al*, 2001) display no major defects in neuronal development, and apparent redundant pathways of neurite outgrowth, mediated by VAMP2, VAMP4 and VAMP7, have been described (Racchetti *et al*, 2010; Schulte *et al*, 2010; Gupton & Gertler, 2010). These evidence raised the possibility that several secretory vesicles equipped with different R-SNAREs, as well as complementary non-vesicular mechanisms, could contribute to neurite extension during brain development. Indeed, we previously found that the R-SNARE Sec22b, a conserved endoplasmic reticulum (ER)-localized R-SNARE involved in vesicle fusion within the early secretory pathway (Xu *et al*, 2000), had an additional and unexpected function in promoting PM expansion during polarized growth. Sec22b concentrates in neuronal growth cones, where it interacts with the neuronal Stx1. Sec22b-Stx1 complex does not mediate fusion, but it rather forms a non-fusogenic bridge between ER and PM. In addition, we showed that increasing the distance between ER and PM, by the insertion of a rigid spacer in Sec22b, reduced neuronal growth, and in budding yeast, the orthologue Sec22/Sso1 complexes contained oxysterol transfer proteins (Petkovic *et al*, 2014). Based on biophysical experiments with synaptic SNAREs (Li *et al*, 2007; Zorman *et al*, 2014), incompletely zippered Sec22b/Stx1 complex would tether ER and PM at distances between 10 and 20 nm, corresponding to the narrowest ER-PM contact sites (Gallo *et al*, 2016).

The critical role of the ER in PM growth is based on its central role in lipid synthesis (Jacquemyn *et al*, 2017). Once synthesized in the ER, lipids travel to the PM along the secretory pathway or they can be directly transferred at ER-PM contact sites. The ER-integral membrane protein Extended-Synaptotagmin (E-Syt) family mediate lipid transfer at ER-PM contact sites. E-Syts are ER-anchored proteins defined by the presence of a cytosolic synaptotagmin-like mitochondrial lipid-binding protein (SMP) domain and multiple Ca^2+^-binding C2 domains (Giordano *et al*, 2013; Min *et al*, 2007). Besides their classical function in tethering ER and PM membranes (Giordano *et al*, 2013), E-Syts transfer lipids via their SMP domains at ER-PM contact sites (Schauder *et al*, 2014; Saheki *et al*, 2016; Reinisch & De Camilli, 2016). On one hand, triple KO of E-Syts 1-3 did not show a major morphological phenotype in neurons suggesting that most function associated with these proteins might be redundant with that of other LTPs (Sclip *et al*, 2016). On the other hand, overexpression of drosophila E-Syt leads to synaptic overgrowth (Kikuma *et al*, 2017) and its knock out led to major growth defect of the mutant flies (Nath *et al*, 2019). Therefore, mammalian E-Syts might be at the same time redundant with other LTPs for bulk growth and limiting factors for specific features of growth and neuronal differentiation. Based on these data, we hypothesized that E-Syts might interact with Sec22b-Stx1 complexes, which in turn could enable bulk ER to PM transfer of lipids responsible for specific features of neurite growth. Here we found a novel interaction between the Sec22b-Stx1 SNARE complex and members of the E-Syt family. We showed that E-Syts were required to stabilize Sec22b-Stx1 association at ER-PM contact sites and that their overexpression in developing neurons promoted axonal growth and ramification which depended on the presence of the SMP and membrane anchoring domains. Furthermore, this E-Syt-mediated morphogenetic effect was inhibited by botulinum neurotoxin C1, which cleaves Stx1 and the expression of Sec22b Longin domain or a mutant with extended SNARE to transmembrane domain linker. These findings support the conclusion that the ternary association between the LTPs E-Syts, Sec22b and Stx plays an important role in plasma membrane expansion leading to axonal growth and ramification.

## Results

### 1. Sec22b and Stx1, 3 interact with the lipid transfer proteins E-Syt2 and E-Syt3

First, we asked whether Sec22b and PM Stx could interact with LTPs. We focused on the E-Syt family of ER-resident LTPs because of their well established presence at ER-PM contact sites and role in glycerophospholipid transfer (Saheki *et al*, 2016; Schauder *et al*, 2014; Yu *et al*, 2016). We performed GFP-trap precipitation experiments on lysates from different cell lines expressing GFP-tagged PM Stx1 and 3 and Sec22b, and tested the presence of E-Syt family members (Fig. 1).

**Figure 1.**
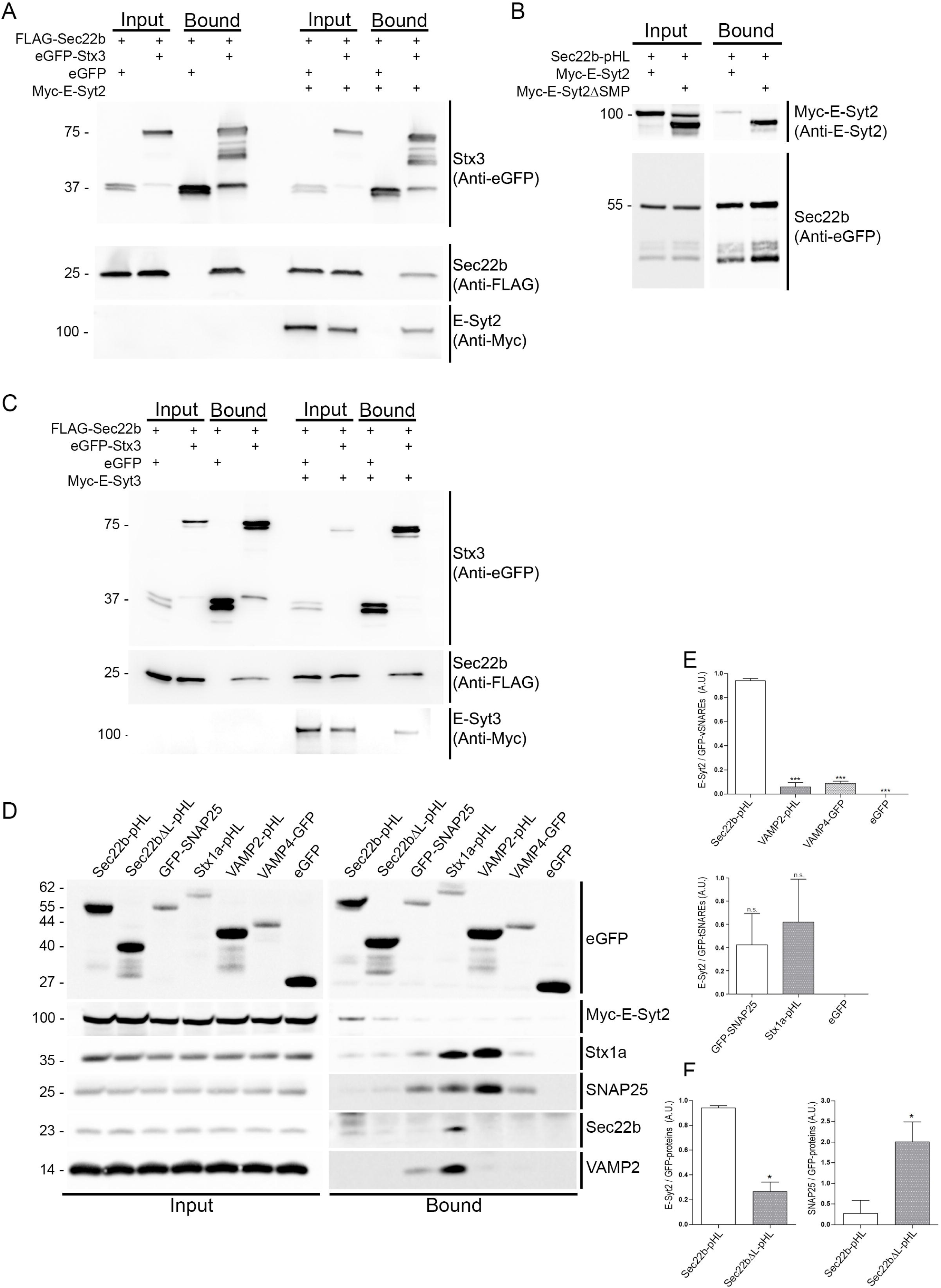
Sec22b and Stx interact with the lipid transfer proteins E-Syt2 and E-Syt3. (A-C) Immunoblots of material recovered after GFP-Trap pull-down from HeLa cell lysates. Cells were transfected for the co-expression of FLAG-Sec22b and Myc-E-Syt2 (A) or FLAG-Sec22b and Myc-E-Syt3 (C) in the presence of either eGFP-Stx3, or eGFP (negative control). Cell lysates were subjected to GFP-Trap pull-down. Total cells lysate (Input) and trapped material (Bound) were processed for SDS-PAGE and Western blotting. Blots were probed with antibodies directed against the tags (GFP, FLAG and Myc) as indicated. Both Myc-E-Syt2 and Myc-E-Syt3 were selectively recruited by eGFP-Stx3. (B) GFP-Trap using Sec22b-pHL as a bait in cells co-expressing either Myc-E-Syt2 or Myc-E-Syt2ΔSMP. Matrix-bound material was processed as in A, B with the indicated antibodies. (D) Immunoblots of material recovered after GFP-Trap pull-down from NGF-treated PC12 cell lysates. Cells were transfected for co-expression of Myc-E-Syt2 and the indicated GFP-tagged SNAREs, and processed as in A-C. Blots were revealed with antibodies against the indicated six target proteins. Only Sec22b-pHL, but not its longin-deleted version Sec22bΔL-pHL or the other tested SNAREs, could recruit Myc-E-Syt2. GFP-Trap pull-down of eGFP was used as control for nonspecific binding. (E) Quantification of the ratio between Myc-E-Syt2 signal and the GFP signal given by the immunoprecipitated GFP-tagged vSNAREs (upper graph) and tSNAREs (lower graph). (F) Quantification of the ratio between Myc-E-Syt2 signal and the GFP signal (left graph) and between endogenous SNAP25 signal and the GFP signal (right graph) given by the immunoprecipitated Sec22b-pHL vs Sec22bΔL-pHL. One-way ANOVA followed by Dunnett’s multiple comparison post-test, P<0,05*, P<0,001***,. n= 3 independent experiments. Error bars represent the SEM.

HeLa cells were used (Fig. 1A-C), which lack neuronal Stx1 but express the closely related homologue Stx3 (Bennett *et al*, 1993). Cells were transfected with the GFP –tagged Stx3 (eGFP-Stx3), together with FLAG-Sec22b and either Myc-E-Syt2 or Myc-E-Syt3, and cell lysates were subjected to GFP-trap. GFP-Stx3 was able to bring down FLAG-Sec22b, further extending previous results obtained with Stx1 (Petkovic *et al*, 2014) (Fig. 1A). A mirror trap experiment using pHLuorin (pHL)-tagged Sec22b (Fig. 1B) comparing the association of full length E-Syt2 and SMP domain-lacking E-Syt2, shows that removal of the SMP domain did not impair binding to Sec22b of E-Syt2 thus suggesting that Sec22b might interact with the C-terminus part of E-Syts. eGFP-Stx3 was also able to precipitate Myc-E-Syt2 (Fig. 1A) as well as Myc-E-Syt3 (Fig. 1C). We then tested whether a Sec22b-Stx-E-Syts association also occurred in a neuronal-like context. PC12 cells were thus chosen because they express neuronal Stx1 and, when treated with NGF, they extend processes similar to those produced by cultured neurons (Greene & Tischler, 1976). Lysates of NGF-differentiated PC12 cells transfected with Myc-E-Syt2 and Sec22b-pHL were subjected to GFP-trap precipitation. Sec22b-pHL co-immunoprecipitated Myc-E-Syt2 (Fig. 1D, first lane). In addition, Sec22b-pHL precipitated small amounts of endogenous Sec22b, and Stx1. To assess the extent of the specificity of the interaction between E-Syt2 and Sec22b, we performed GFP-trap in lysates of NGF-differentiated PC12 cells co-expressing Myc-E-Syt2 and various pHL- or GFP-tagged v- and t-SNAREs, i.e. Sec22bΔL-pHL, the mutant Sec22b lacking the N-terminal Longin domain, GFP-SNAP25, Stx1-pHL, VAMP2-pHL, VAMP4-GFP, and GFP alone as indicated (Fig. 1D). A trace amount of E-Syt2 was found to co-precipitate with GFP and reduced amounts were found in the precipitates of GFP-SNAP25, Stx1-pHL, VAMP2-pHL, VAMP4-GFP (Fig. 1E). The amount of co-precipitated E-Syt2 was greatly reduced while the amount of SNAP-25 became clear in the case of Sec22bΔL-pHL mutant, as compared to wild-type Sec22b (Fig. 1F). These data strongly suggest that Sec22b association with E-Syt2 is specific and requires the N-terminal Longin domain of Sec22b. These results are compatible with Longin domain-dependent presence of Stx1 in Sec22b/E-Syt2 complexes. The fact that Sec22bΔL-pHL precipitated SNAP-25 whereas it was barely detectable in the case of Sec22b-pHL suggests that the Longin domain may prevent SNAP-25 from entering the Stx1/Sec22b complex. It is interesting to note, as positive control, that Stx1-pHL and GFP-SNAP-25 co-precipitated large amounts their endogenous synaptic SNARE complex partners (Stx1, SNAP-25, VAMP2).

Taken together GFP-trap experiments support the notion that Sec22b-PM Stx complexes associate with members of the E-Syt family of LTPs and that this interaction occurs both in a non-neuronal and neuronal cellular context. Comparison with the typical synaptic SNARE complex suggests that Stx1 would associate primarily with SNAP-25 and VAMP2 whereas complexes with Sec22b would represent a minor pool of Stx1, in agreement with their restricted occurrence of ER-PM contact sites.

To go further into the characterization of the Sec22bΔL mutant, we used surface staining to detect the presence of wild type or mutant Sec22b at the plasma membrane with ectoplasmic antibodies added to living cells. We used again Sec22b-pHL and Sec22bΔL-pHL chimera in which Sec22b and its mutant version are C-terminally-tagged with pHL, so that anti-GFP antibody present in the medium would bind and reveal only Sec22b that reached the cell surface. N-terminally tagged Sec22b served as a negative control in these experiments, since its GFP tag should never be exposed extracellularly thus be undetectable with anti-GFP antibody without permeabilization. As a positive control for fusion, we used VAMP2-pHL (Fig. S1A). In contrast to the strong positive signal given by VAMP2, we found no detectable Sec22b at the plasma membrane, consistent with its non-fusogenic function at the PM (Petkovic *et al*, 2014) (Fig. S1B). On the contrary, significant amounts of Sec22bΔL were detected at the cell surface (Fig. S1B). Thus, Sec22bΔL was able to mediate fusion to a certain degree with the plasma membrane. In addition, in COS7 cells, Sec22bΔL mutants did not display the ER localization typical of Sec22b wild type, but it was also found in vesicular-like structures (Fig. S1C), leading to the hypothesis that it might in part escape into secretory vesicles that eventually fuse with the PM because Sec22bΔL was able to bind both Stx1 and SNAP-25 (Fig. 1D) These data further suggested the involvement of the Longin domain of Sec22b in the formation of the non-fusogenic Sec22b/Stx1 complex at ER-PM contact sites.

### 2. E-Syt2 and Sec22b are in close proximity to the plasma membrane in neurites and growth cones

To determine whether the observed association between Sec22b and E-Syts could also be observed in cells *in situ*, we used *in situ* Proximity Ligation Assay (PLA), live cell imaging and and Stimulated Emission Depletion (STED) super-resolution microscopy.

Fixed HeLa cells and hippocampal neurons at 3DIV were stained with Sec22b and E-Syt2 antibodies and processed for PLA. As specificity control of the PLA signal given by Sec22b and E-Syt2, we also tested the proximity of Sec22b with Calnexin, another ER-resident protein not supposed to interact with Sec22b (Fig. 2). Distinct PLA dots were observed in HeLa cells labeled for endogenous Sec22b and E-Syt2 (7.18 ± 0.60 per cell) (Fig. 2A). The PLA signal decreased to an average of 3.30 ± 0.40 in cells stained for Sec22b and Calnexin and to an average of 1.15 ± 0.14 and 1.20 ± 0.11 in negative controls, performed by incubating cells with only Sec22b and E-Syt2 antibodies, respectively (Fig. 2B). To further confirm these results, we performed live cell imaging in HeLa cells co-expressing mCherry-Sec22b and GFP-E-Syt2. We found co-localization in discrete puncta of Sec22b and E-Syt2, prominently localized in the periphery of the cell. Interestingly, such puncta appeared to be less dynamic compared to corresponding whole protein populations, suggesting that they could represent hot-spots of membrane contact sites populated by Sec22b and E-Syt2 (Movie 1). In neurons, the amount of PLA signal per cell was normalized for the area of neurites or cell body (Fig. 2C). Interestingly, dots revealing Sec22b-E-Syt2 proximity were more numerous in neurites (5.53% ± 0.60%) (Fig. 2D) and growth cones (7.64% ± 1.49) (Fig.2E) compared to cell bodies (2.85% ± 0.45%) (Fig. 2F), suggesting a preferential interaction of the two proteins in growing processes. Similarly, the PLA signal between Sec22b and Calnexin and in the negative control was strongly reduced as compared to that of Sec22b-E-Syt2 in neurons.

**Figure 2.**
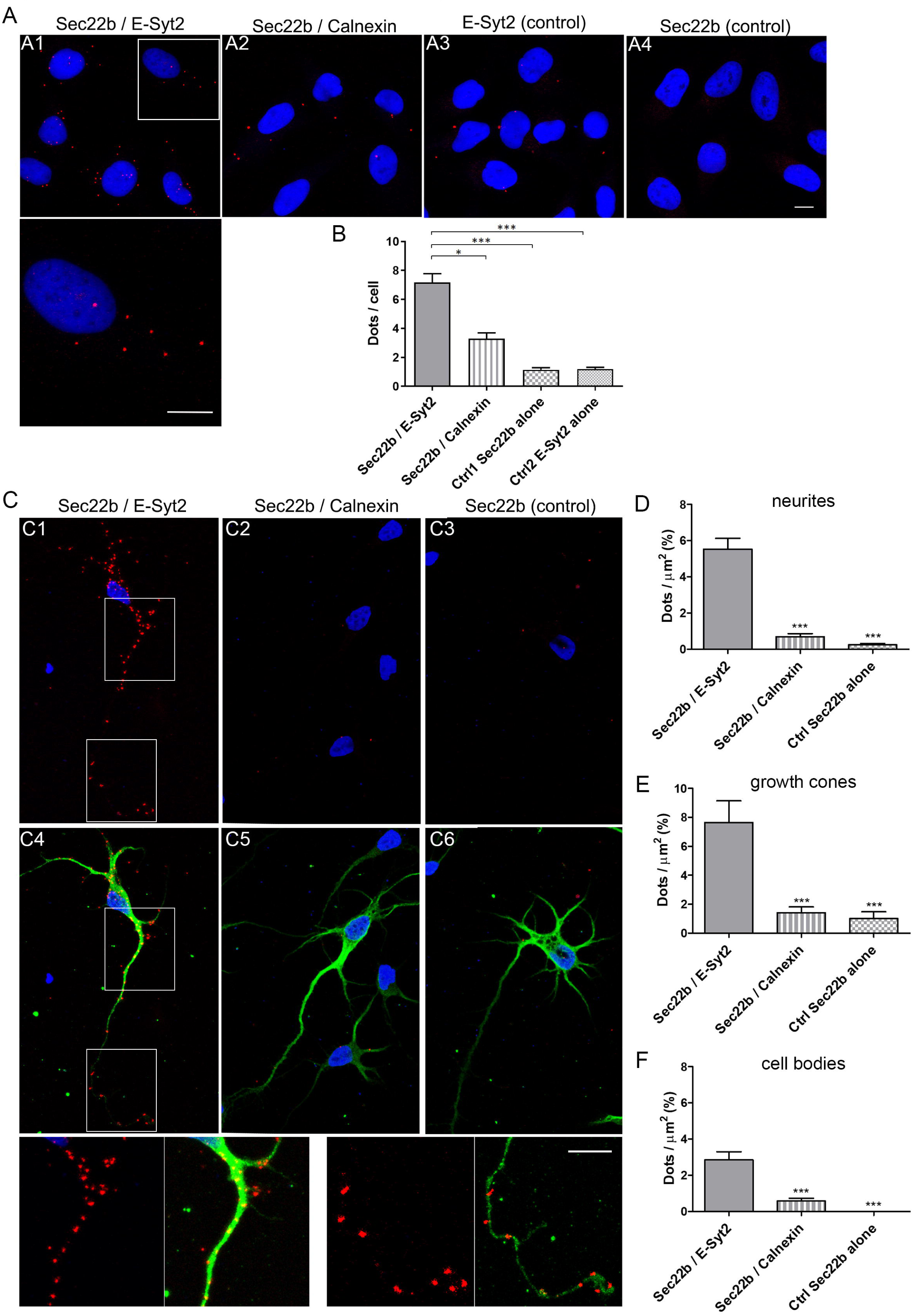
E-Syt2 and Sec22b are abundantly in close proximity in neurites and growth cones. Duolink proximity ligation assay (PLA) for protein interactions *in situ* was performed in HeLa cells (A) and 3DIV hippocampal neurons (C). Representative confocal images are shown for the indicated antibody combinations using mouse anti-Sec22b, rabbit anti-E-Syt2, or rabbit anti-Calnexin. Negative controls consisted in using anti-Sec22b or anti-E-Syt2 antibody only. In each field, maximum intensity projection of a confocal z-stacks including a whole cell were performed to observe the maximum amount of PLA dots (red). Nuclei were stained by DAPI (blue). A1-4, PLA dots. Lower panel is a higher magnification of region outlined in A1. C1-3, PLA dots. C4-6, MAP2 immunofluorescence staining superimposed to fields shown in C1-3. Lower panels are a higher magnifications of region outlined in C1 and C4. Scale bar, 10 µm. (B, D, E) Quantification of PLA is expressed as dots per HeLa cell (B), and in 3DIV hippocampal neurons as dots per µm^2^ of surface area in neurites (D), growth cones (E) or in cell body (F). The number of individual fluorescent dots is higher in the Sec22b-E-Syt2 pair as compared to Sec22b-Calnexin pair or negative controls both in HeLa and in neurons. It is higher in neurites and growth cones as compared to cell bodies in neurons. One-way ANOVA followed by Dunn’s multiple comparison post-test, P<0,05*, P < 0.001***. n=3 independent experiments. Error bars represent the SEM.

Close proximity between Sec22b-E-Syt2 was confirmed by 3D STED microcopy visualizing Sec22b-pHL and endogenous E-Syt2 localization in growth cones at high resolution (Fig. 3A, B). Spatial distribution analysis of E-Syt2-and Sec22 was done using “Icy SODA” plugin and Ripley’s function (Lagache *et al*, 2018). When statistically associated, both Sec22b and E-Syt2 were detected in growth cones at an average distance of 84.33 nm ± 9.64 nm of each other. Shorter distances between the two proteins were measured in all the analyzed growth cones. Furthermore, we measured the distance between these two molecules and the PM (Fig. 3C). To this end, we labelled the PM with Wheat Germ Agglutinin (WGA), which allows the detection of glycoconjugates, *via* N-acetylglucosamine and N-acetylneuraminic acid (sialic acid) residues, on cell membranes. We found that median distance d (Figure 3C) between plasma membrane and E-Syt2 coupled to Sec22b-pHL was 33.6 nm (4 independents growth cones, 4226 clusters) suggesting that when E-Syt2 and Sec22b are associated they populate ER-PM contact sites to a large extent.

**Figure 3.**
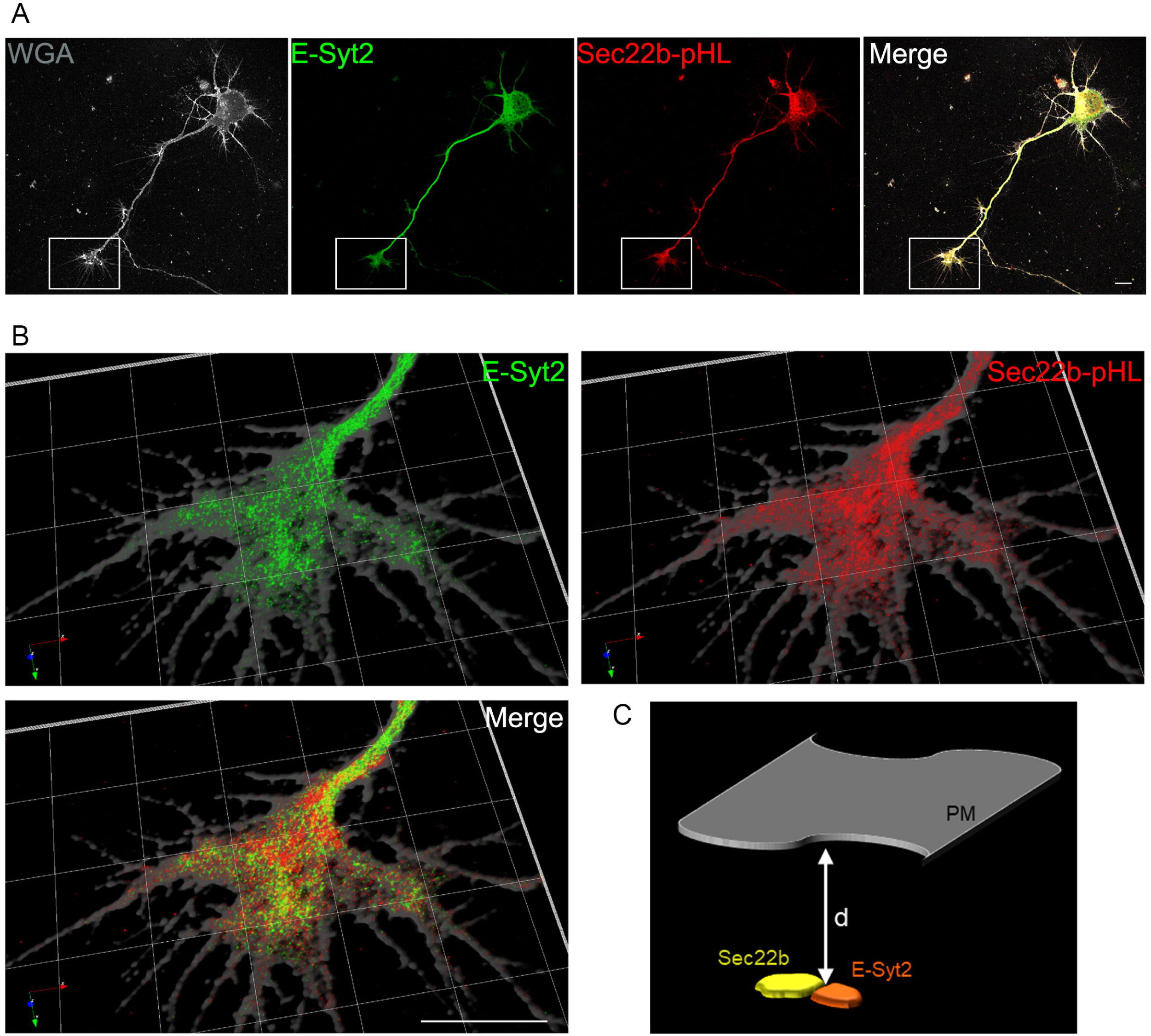
Analysis of E-Syt2 / Sec22b association using super-resolution microscopy. (A) Representative confocal images of a 3DIV hippocampal neuron labeled for endogenous E-Syt2 (green) and Sec22b-pHL (red) and plasma membrane (gray). Alexa Fluor® 488- conjugated Wheat Germ Agglutinin (WGA) was used to label the plasma membrane (grey). Scale bar, 10 μm. (B) Super-resolution (3D STED) 3D reconstruction of the inset in (A) showing localization of endogenous E-Syt2 and Sec22b-pHL in the growth cone. (C) Statistical association was assessed through spatial distribution analysis using Ripley’s function (Icy SODA Plugin). Topological scheme illustrating the measured distance (d) between associated Sec22b-E-Syt2 puncta and the PM. d was estimated to an average of 67.12nm+/-1.22 in four different growth cones for 4229 clusters and its median was: 33.6nm indicating that 50% of the clusters were at a distance ≤d=33.6nm, compatible with ER-PM contact sites.

### 3. E-Syts favor the occurrence of Sec22b-Stx complexes

As previously reported, E-Syts function as regulated tethers between the ER and the PM and their overexpression stabilizes and increases the density of ER-PM contact sites (Giordano *et al*, 2013). The question as to whether E-Syts overexpression stabilizes Sec22b-Stx complexes at MCSs was therefore addressed. To do so the PLA technique was used to evaluate Sec22b/Stx3 association in non-transfected or in E-Syt3-overexpressing HeLa cells (Fig. 4A, B). We found that, the number of PLA puncta significantly increased from 4.08 ± 0.67 per cell dots counted in non-transfected cells to 6.66 ± 0.91 in cells where E-Syt3 is overexpressed (Fig. 4C). We then investigated to which extent impairing the formation of ER-PM contact sites would influence the interaction between Sec22b and Stx3. To do this, we inhibited expression of the three E-Syt isoforms (E-Syt1, E-Syt2 and E-Syt3) simultaneously using siRNAs targeting the mRNAs of these proteins (Fig. 4D, E). Interestingly, the total number of PLA dots given by Sec22b and Stx3 was strongly decreased in the E-Syt-deficient HeLa cells as compared to control cells (Fig. 4F).

**Figure 4.**
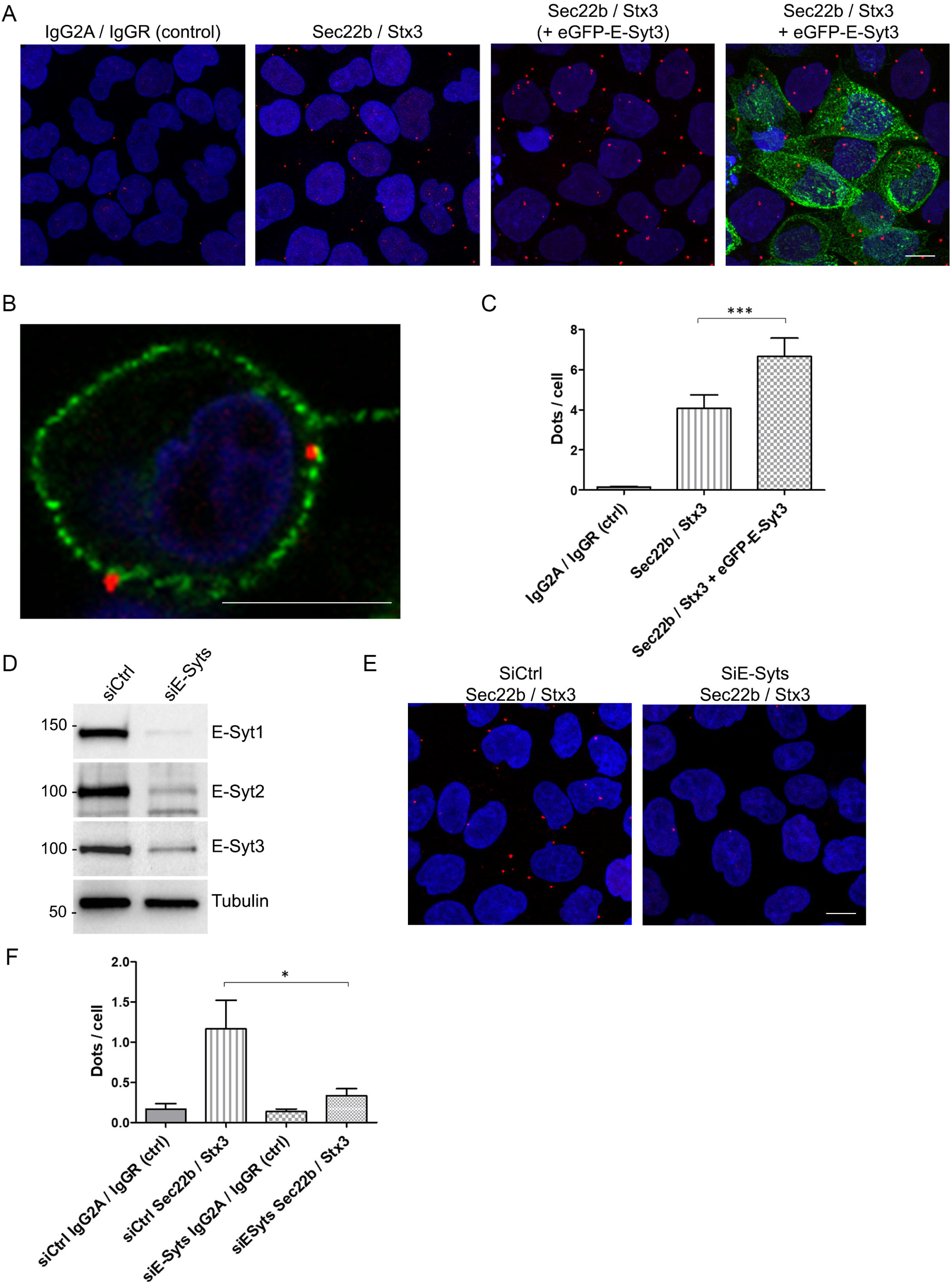
E-Syts favor the occurrence of Sec22b-Stx complexes. Duolink proximity ligation assay (PLA) for protein interactions *in situ* was performed in HeLa cells either non transfected or overexpressing eGFP-E-Syt3 (A) and in HeLa cells expressing siRNAs targeting E-Syt1, E-Syt2 and E-Syt3 simultaneously (E). Representative confocal images are shown for PLA between Sec22b (Mouse anti-Sec22b) and Stx3 (Rabbit anti-Stx3). Negative controls combined IgG2a and IgGR. In each field, maximum intensity projection of a confocal z-stacks including a whole cell were performed to observe the maximum amount of PLA dots (red). Nuclei were stained by DAPI (blue). Scale bar, 10 µm. (B) Confocal maximal projection image showing co-localization of Sec22-Stx3 PLA signal and E-Syt3-positive cortical ER. Scale bar, 10 μm. (C, F) Quantification of PLA is expressed as dots per HeLa cell. The number of individual fluorescent dots is higher in cells overexpressing eGFP-E-Syt3 as compared to non-transfected cells or negative control (C). It is lower in cells expressing siRNAs targeting the three E-Sys isoforms as as compared to cells expressing siCtrl (F). Data are expressed as means ± SEM. Student t-test P<0,05*, P<0,001***. n= 3 independent experiments. (D) Representative immunoblots from lysates of HeLa cells expressing siRNAs targeting the three E-Sys isoforms. Tubulin was used as a loading control.

Taken together, our results indicate that increasing the amount of or abolishing the tethering activity of E-Syts correlates with the probability to observe proximity between Sec22b and Stx3. Thus, E-Syts likely promote the formation of ER-PM contact sites populated by Sec22b-Stx3 complexes.

### 4. E-Syts overexpression promotes filopodia formation and ramifications in developing neurons

At the neuromuscular junction of Drosophila melanogaster, overexpression of *Esyt* orthologue, enhances synaptic growth (Kikuma *et al*, 2017). This prompted us to test the effect of E-Syts overexpression in the growth of cultured neurons. Therefore, rat hippocampal neurons were nucleofected with Myc-E-Syt2 and cultured until 3DIV, a time when axonal polarization is achieved and the major axonal process is distinguishable from the minor dendritic neurites (Dotti *et al*, 1988). eGFP co-transfection was included to appreciate neuronal morphology of transfected neurons. Overexpression of E-Syt2 induced the formation of actin-positive filopodia and ramifications, particularly in the growing axons (Fig. 5A; Fig. S2). Comparison of the phenotype of neurons expressing Myc-E-Syt2, and neurons expressing E-Syt2 mutant versions, lacking either the SMP (Myc-E-Syt2 ΔSMP[119-294]) or the membrane spanning domain (Myc E-Syt2 ΔMSD[1-72]) domains was undertaken using the Myc-Empty vector as negative control (Fig. 5B). Quantification of morphological parameters of transfected cells showed that neurons overexpressing Myc-E-Syt2 displayed a ∼1.5- and a ∼2-fold increase in the total neurite length and in the number of branching, respectively, as compared to E-Syt2 mutants or negative control (Fig. 5C). Surprisingly, despite a tendency measured as a ∼35% increase, the major neurite was not significantly longer in Myc-E-Syt2 overexpressing neurons (Fig. 5C). Noteworthy, E-Syt2 mutants exhibited no significant differences in the parameters analyzed when compared to the negative control. Thus, elevating the expression level of wild type but not mutant E-Syt2 in developing neurons clearly promoted neurite growth and ramification.

**Figure 5.**
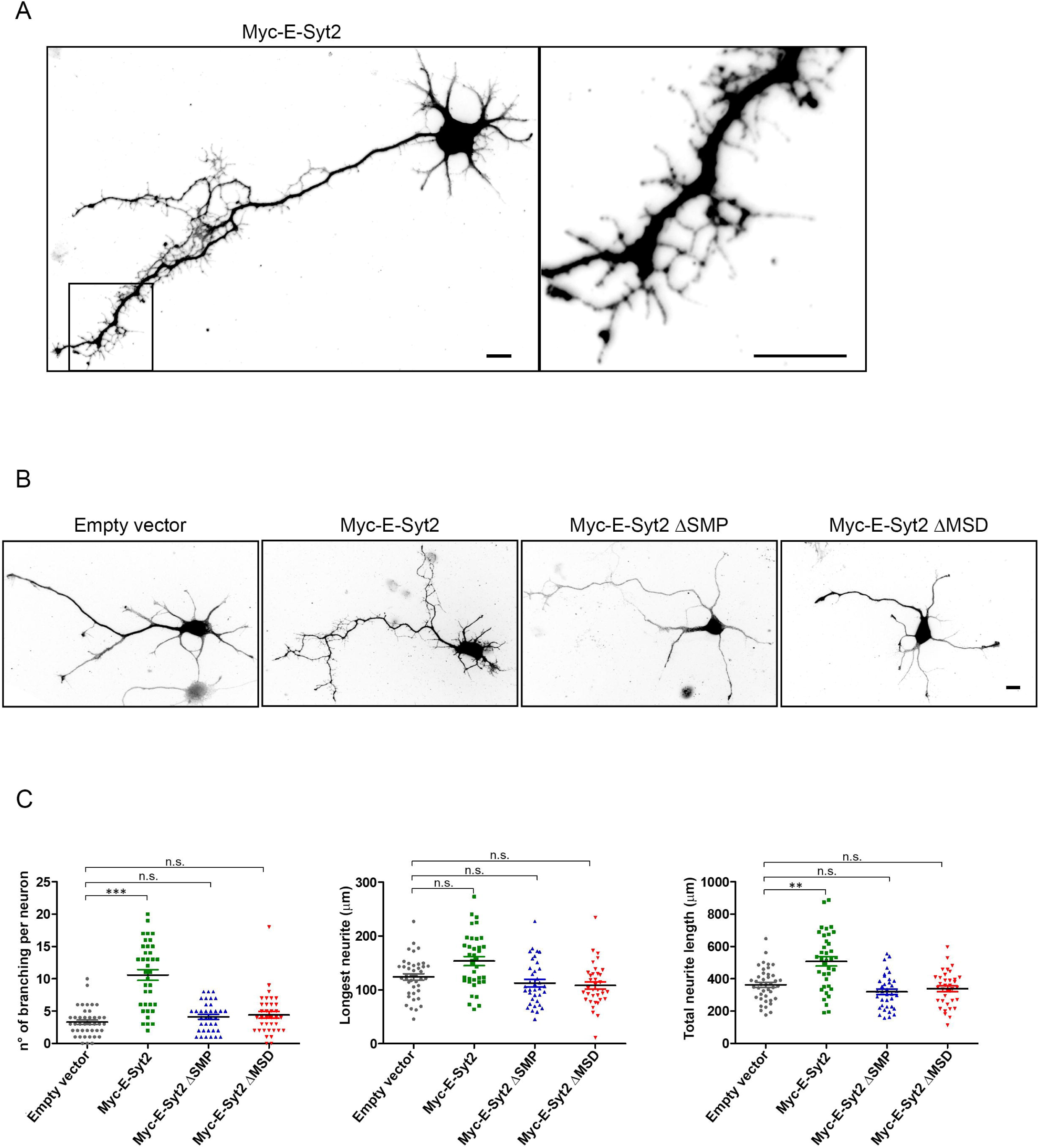
E-Syts overexpression promotes filopodia formation and ramifications in developing neurons. (A) Myc-E-Syt2 expression in 3DIV hippocampal neurons. A2, representative morphology of a nucleofected neuron expressing Myc-E-Syt2. eGFP co-expression was used to view cell shape. A2, higher-magnification of inset in A1 showing high density of filopodia in a segment of the axonal shaft. Scale bar, 10 µm. (B) Expression of Myc-E-Syt2 mutated forms. 3DIV neurons co-expressing eGFP and Myc-E-Syt2 or one of the deletion mutants Myc-E-Syt2ΔSMP and Myc-E-Syt2ΔMSD, lacking the SMP domain [119-294] and the membrane spanning region [1-72], respectively. The empty Myc vector was used as negative control. Scale bar, 10 µm. (C) Quantification of morphological parameters in transfected neurons shown in B. Plots were acquired on maximal intensity projections of z-stacks of the eGFP channel. Note that in comparison with the longest neurite length (C2), the number of branching per neuron (C1) and the total neurite length (C3) are increased in neurons upon Myc-E-Syt2 expression, whereas expression mutants had no effect. Data are expressed as means ± SEM. Oneway ANOVA P<0,001***, Dunn’s multiple comparison post-test labeled on graph. n= 3 independent experiments.

### 5. E-Syts overexpression stimulates membrane growth in HeLa cells

To assess whether the increase in growth observed in neurons is a general feature resulting from E-Syt2 overexpression, we compared the phenotype elicited by expression of Myc-E-Syt2 and its deleted mutants in non-neuronal HeLa cells, following an experimental paradigm similar to that used in the previous experiment. Briefly, we transfected HeLa with Myc-E-Syt2, Myc-E-Syt2 ΔSMP[119-294], Myc E-Syt2 ΔMSD[1-72] or Myc-Empty vector together with eGFP. Two days after transfection, cell were fixed and the eGFP immunostaining was used to allow a phenotype comparison in the different conditions (Fig. 6A, B). Consistent with the observations in developing neurons, a clear morphogenetic effect was observed. Myc-E-Syt2 overexpressing HeLa cells displayed an enhanced filopodia formation as compared to control and mutants overexpression, as measured by the percentage of the PM spikes area over the total cell surface (Fig. 6C). Taken together, these results provide additional evidence of the involvement of E-Syts in membrane growth, both in neuronal and non-neuronal contexts.

**Figure 6.**
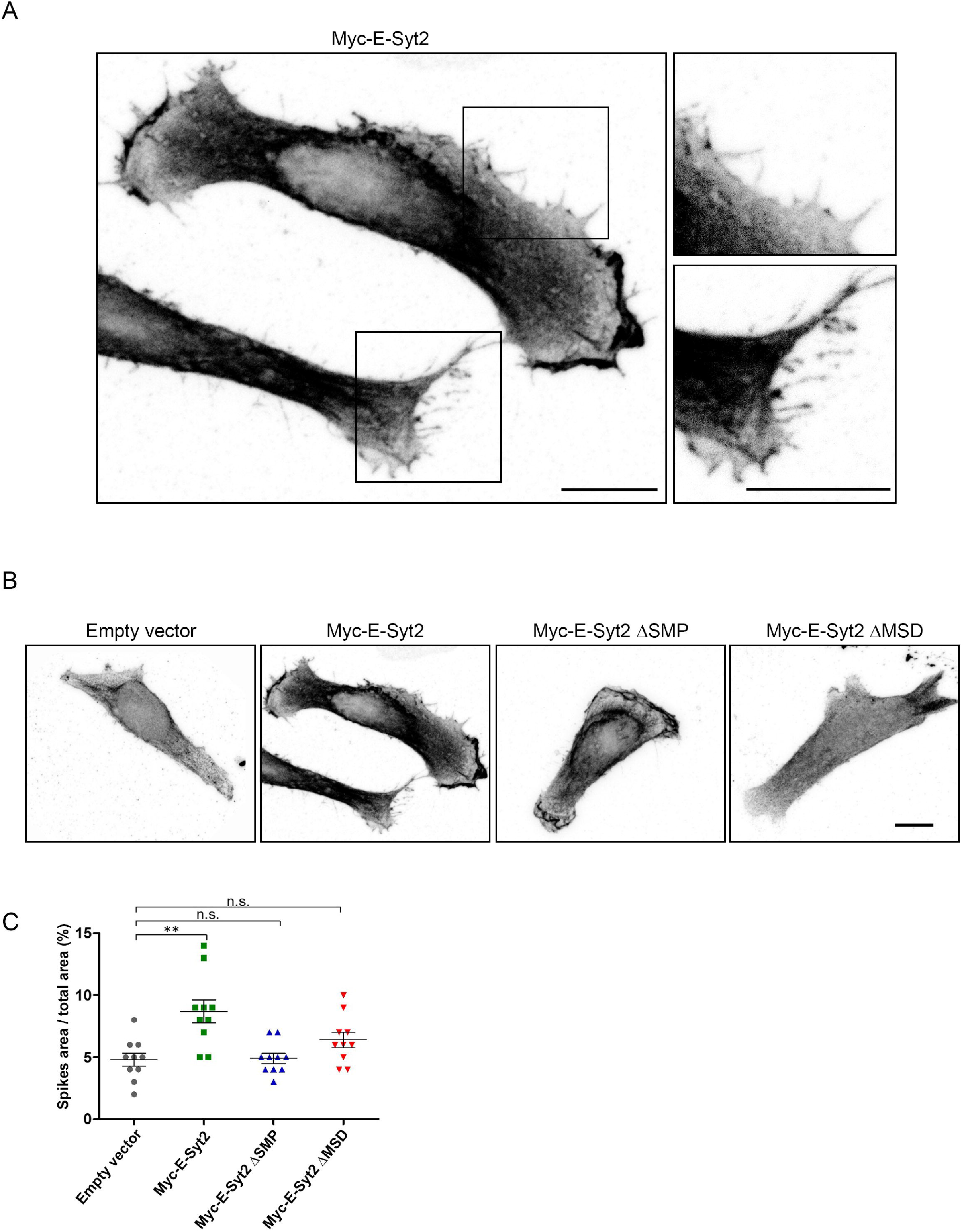
E-Syts overexpression stimulates membrane growth in HeLa cells. (A) HeLa cells were transfected for co-expression of Myc-E-Syt2 and eGFP (the latter to delineate cell shape). Shown is a representative image of two cells (A1). A2, 3, higher magnifications of regions outlined in A1, showing high density of filopodia at the cell periphery. Scale bar, 10 µm. (B) HeLa cells co-expressing eGFP and Myc-E-Syt2 or one of the deletion mutants Myc-E-Syt2ΔSMP and Myc-E-Syt2ΔMSD as in figure 2. The empty Myc vector was used as control. Scale bar, 10 µm. (C) Quantification of spike area in transfected cells shown in B, acquired from maximal intensity projections of z-stacks of the eGFP channel. It is expressed as percentage of the total cell surface area. Filopodia formation was enhanced in cells expressing Myc-E-Syt2 as compared to cells expressing the mutant proteins. Data are expressed as means ± SEM. Oneway ANOVA P<0,01**, Dunn’s multiple comparison post-test labeled on graph. n= 3 independent experiments.

### 6. E-Syts-mediated morphogenetic effect depends on Stx1

Next, the role of the ternary assembly of E-Syt2, Sec22b and Stx1 in the phenotype of enhanced neuronal growth elicited by E-Syt2 was investigated. To this end, we designed experiments aimed at impairing Stx1 in neurons overexpressing Myc-E-Syt2. Enzymatic disruption of the SNARE Sec22b-Stx1 complex was achieved using botulinum (BoNT) neurotoxins, zinc-endoproteases known to cleave SNARE proteins (Binz *et al*, 2010; Binz, 2013; Sikorra *et al*, 2008). For this study, purified BoNT/A was used to cleave SNAP25, BoNT/C1 to cleave both Stx1 and SNAP25, and BoNT/D to cleave VAMP2 (Fig. 7A). We nucleofected rat hippocampal neurons with Myc-E-Syt2 and, before fixation at 3DIV, we incubated cells with BoNTs (as indicated in figure legend). Cells incubated with the toxins-diluting culture medium were used as negative control. Since exposure to BoNT/C1 at high concentrations and for long incubation periods causes degeneration of neurons in culture (Osen-Sand *et al*, 1996; Igarashi *et al*, 1996), various concentrations and incubation times were tested, and a 4-hour treatment of neurons with 1nM BoNTs was chosen to avoid such deleterious effects. This low concentration of toxins, associated with a relatively short incubation period, is sufficient to induce SNARE cleavage, as shown by reduced protein signals detected after Western blotting (Fig. 7B, C). Assuming that Sec22b, Stx1 and E-Syt2 form a complex, only BoNT/C1 was expected to prevent the E-Syt2-induced phenotype of growth, since this would occur following a cleavage of Stx1. BoNT/A and D, acting on SNAP25 and VAMP2 respectively, should have no effect. Noticeably, the cleavage of Stx1, occurring after BoNTC/1 incubation, was found to cause a ∼50% decrease in the extent of branching, resulting in a ∼20% reduction of the total neurite length, as compared to neurons incubated with the other tested BoNTs and negative control (Fig. 7D, E). No significant decrease in the length of the major neurite was observed in cells after BoNT/C1 incubation.

**Figure 7.**
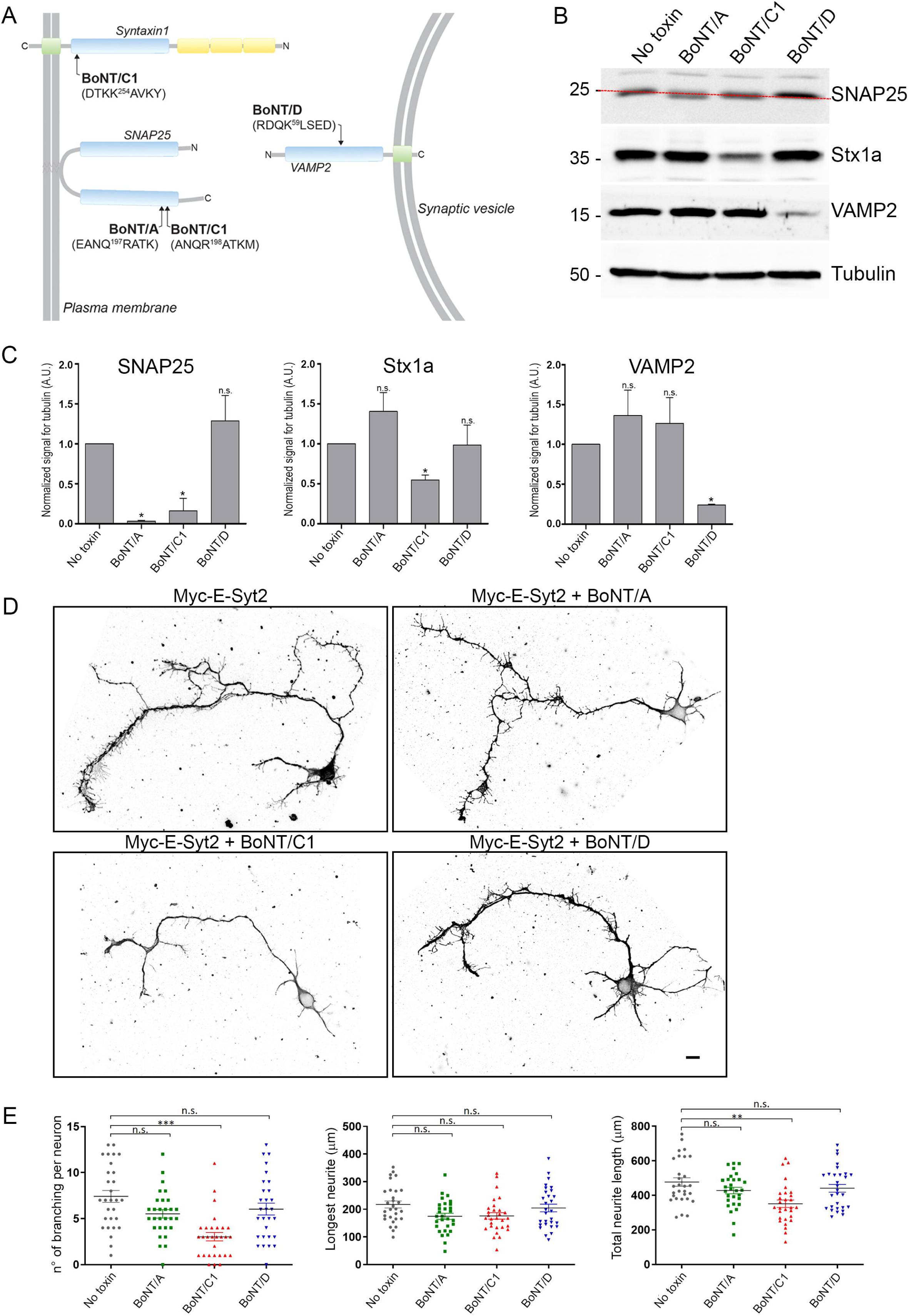
E-Syts-mediated morphogenetic effect depends on Stx1. (A) Schematic of cleavage sites on neuronal SNARE targets of BoNTs. Cleavage can occur on SNAP25 (BoNT/A), on both SNAP25 and Stx1 (BoNT/C1), or specifically on VAMP2 (BoNT/D). (B1, 2) Cleavage activity of BoNTs. (B1) Representative immunoblots from lysates of neurons exposed for 4 h to 1 nM BoNTs. (B2) Quantification of ECL signals from B1. Ratios of SNAREs to tubulin are plotted. Oneway ANOVA P<0,05*, followed by post-hoc Tukey HSD test. n=3 independent experiments. Data are expressed as means ± SEM. (D) 3DIV hippocampal neurons co-expressing Myc-E-Syt2 and eGFP and treated with BoNTs (1 nM, 4h). Scale bar, 10 µm. (E) Quantification of morphometric parameters on treated neurons, measured on maximal intensity projections of z-stacks of the eGFP channel. The specific cleavage of Stx1 by BoNT/C1 reduces the number of ramifications and the total neurite length of Myc-E-Syt2 expressing neurons. Data are expressed as means ± SEM. Oneway ANOVA P<0,01** P<0,001 ***, Dunn’s multiple comparison post-test labeled on graph. n= 3 independent experiments.

These results support the conclusion that Stx1 is required for E-Syt2 to promote neuronal growth, as particularly stressed out by ramification and filopodia formation.

### 7. E-Syts-mediated morphogenetic effects depend on the close apposition of ER to PM mediated by Sec22b-Stx1 complexes

In view of the data reported above, it was necessary to investigate whether the distance between ER and PM of contact sites formed by Sec22b and Stx1 was a determining factor for the acquisition of the E-Syt2-mediated morphological phenotype. To this end, the GFP-Sec22b-P33 mutant was used as previously described (Petkovic *et al*, 2014). In GFP-Sec22b-P33 the SNARE and transmembrane domains of Sec22b are linked by a stretch of 33 prolines (Fig. 8A). Electron tomography analysis showed that GFP-Sec22b-P33 expression resulted in a 6-nm increase of of the ER to PM distance at contact sites, without changing Sec22b localization and its interaction with Stx1 (Petkovic *et al*, 2014). Co-expression of the GFP-Sec22b-P33 mutant with Myc-E-Syt2 was found to prevent the increase in branching observed in neurons overexpressing E-Syt2 (Fig. 8B, C). This data confirmed and extended results obtained by impairing Stx1 via BoNT/C1. They support the notion that, for E-Syt2 to perform its function in membrane growth, the strict structure of the Sec22b-Stx1 complex is required at ER-PM contact sites.

**Figure 8.**
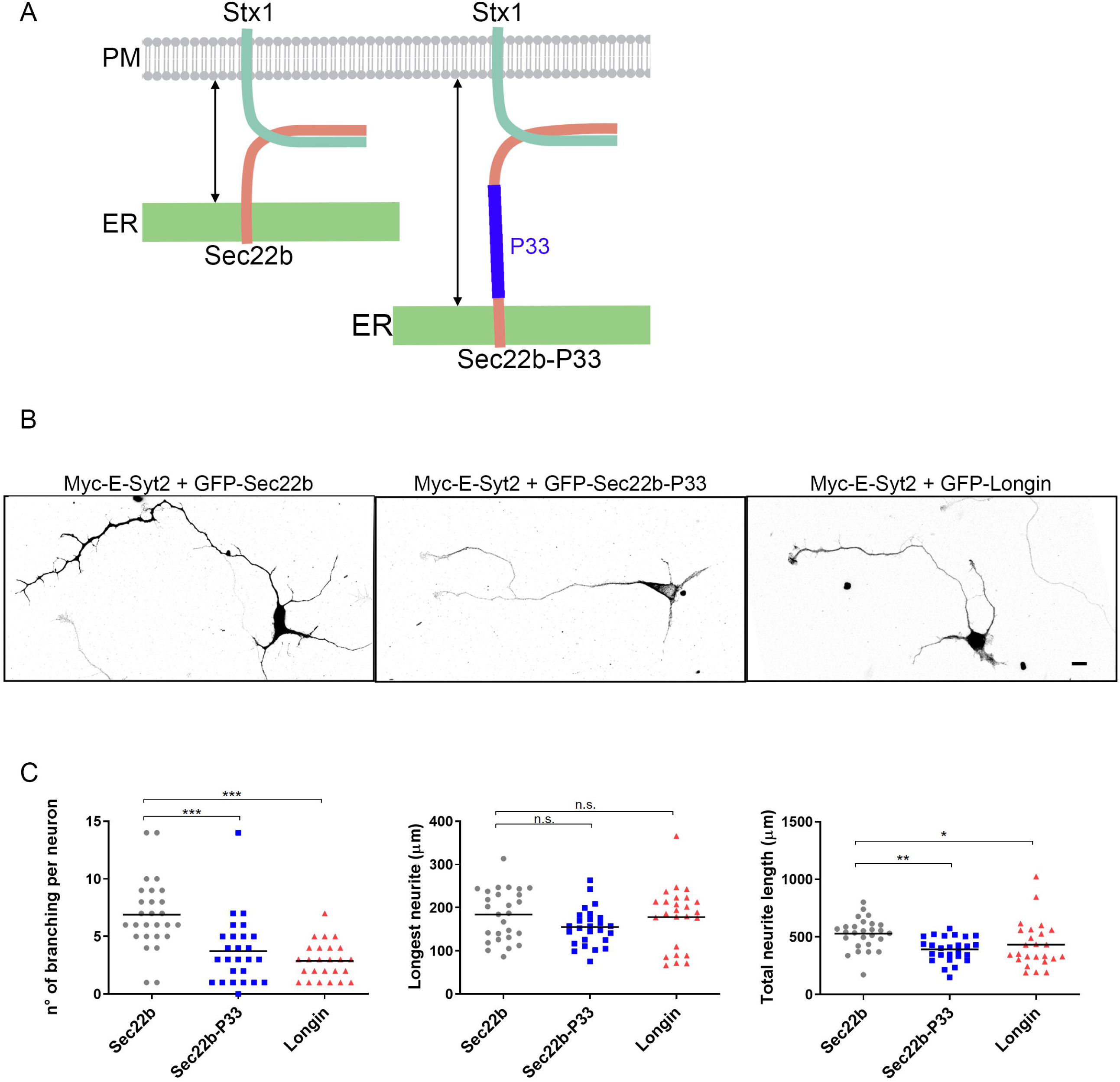
E-Syts-mediated morphogenetic effects depend on ER to PM distance. (A) Scheme showing the predicted effect of a polyproline stretch insertion in Sec22b on the ER-PM distance (see text). (B) Hippocampal neurons co-expressing Myc-E-Syt2 and either GFP-Sec22b, GFP-Sec22b-P33 mutant or Sec22b GFP-Longin domain, observed at 3DIV. Scale bar, 10 µm. (D) Quantification of morphological parameters on treated neurons, measured on maximal intensity projections of z-stacks of the GFP channel. Effect of overexpressed Myc-Esyt2 on number of ramifications and on the sum of neurite lengths is reduced in neurons expressing the GFP-Sec22b-P33 mutant or the GFP-Longin domain. Data are expressed as means ± SEM. Oneway ANOVA P<0,01** P<0,001 ***, Dunn’s multiple comparison post-test labeled on graph. n= 3 independent experiments.

To gain further insight in the requirement of Sec22b in E-Syts mediated neurite growth, the effect of co-expressing the Sec22b Longin domain with E-Syt2 was tested because we showed previously (Fig. 1) that the Longin domain was important for Sec22b/E-Syt2 interaction. Similarly to what was previously reported for the Sec22b-P33 mutant, the expression of the Sec22b Longin domain was found to reduce branching and the overall neurite length in Myc-E-Syt2 overexpressing hippocampal neurons (Fig. 8B, C).

## Discussion

In the present study, we report on a novel interaction between two ER-resident membrane proteins, the R-SNARE Sec22b and members of the Extended-Synaptotagmin (E-Syts) family and show the functional relevance of this interaction in neuronal differentiation. Indeed we found 1/ an interaction between Sec22b and E-Syt2 which required the Longin domain of Sec22b, 2/ this interaction occurred at ER-PM contact sites particularly in neurites, 3/overexpression of wild type E-Syt2 but not mutants devoid of lipid transfer or membrane anchoring domains increased filopodia formation, neurite growth and ramification, and 4/ this morphological effect of E-Syt2 overexpression required functional Stx1 and Sec22b.

### LTPs as partners of SNARE complexes at MCSs

GFP-trap pull down experiments using Sec22b as a bait systematically captured E-Syts. The N-terminal Longin domain of Sec22b had a prominent role in this interaction because the Longin deletion mutant of Sec22b had a reduced ability to capture E-Syts. PM Stx1 and Stx3 also precipitated Sec22b and E-Syts suggesting a ternary complex. This complex however likely corresponded to a small pool of Stx1 in PC12 cells, when compared to the synaptic SNARE complex. Notably, SNAP25 coprecipitated with Sec22b at a level just above detection and deletion of the Longin domain increased the amount of coprecipitated SNAP25. This is consistent with the lack of SNAP-23/25/29/47 in Stx1/Sec22b complexes (Petkovic *et al*, 2014) and the notion that the Longin domain may play a role to exclude SNAP25. Moreover, the 1:1 complex Stx1 and SNAP25 is very abundant and it precedes the recruitment of VAMP2 leading to the formation of the ternary synaptic SNARE complex essential for neurotransmitter release (Fasshauer & Margittai, 2004; Weninger *et al*, 2008). Hence, we are led to propose a model whereby the binding of a pre-assembled, ER-resident, Sec22b/E-Syts complex to a PM-resident Stx1/SNAP25 complex might displace SNAP25 from Stx1 leading to the formation of a non-fusogenic ternary assembly of Sec22b, Stx1 and E-Syts and this hypothetical sequence of event would depend on the presence of the Longin domain (Fig. 9). Other LTPs are expected to interact with Sec22b and PM Stx as already in yeast we previously found Osh2/3 associated with Sso1 (Petkovic *et al*, 2014). Furthermore, in mammalian cells, both Sec22b and Stx1 were shown to interact with the ER-resident VAMP-associated protein VAP-A. (Weir *et al*, 2001). VAP-A mediates stable ER-PM tethering (Loewen *et al*, 2003) and binds a wide number of LTPs, such as oxysterol-binding protein (OSBP)-related proteins (ORPs) and ceramide transfer protein (CERT) (Lev, 2010). Identifying the full catalogue of LTP associated with PM Stx and Sec22b will thus be an important future direction to understand in details the molecular mechanisms occurring at ER-PM contact sites.

**Figure 9.**
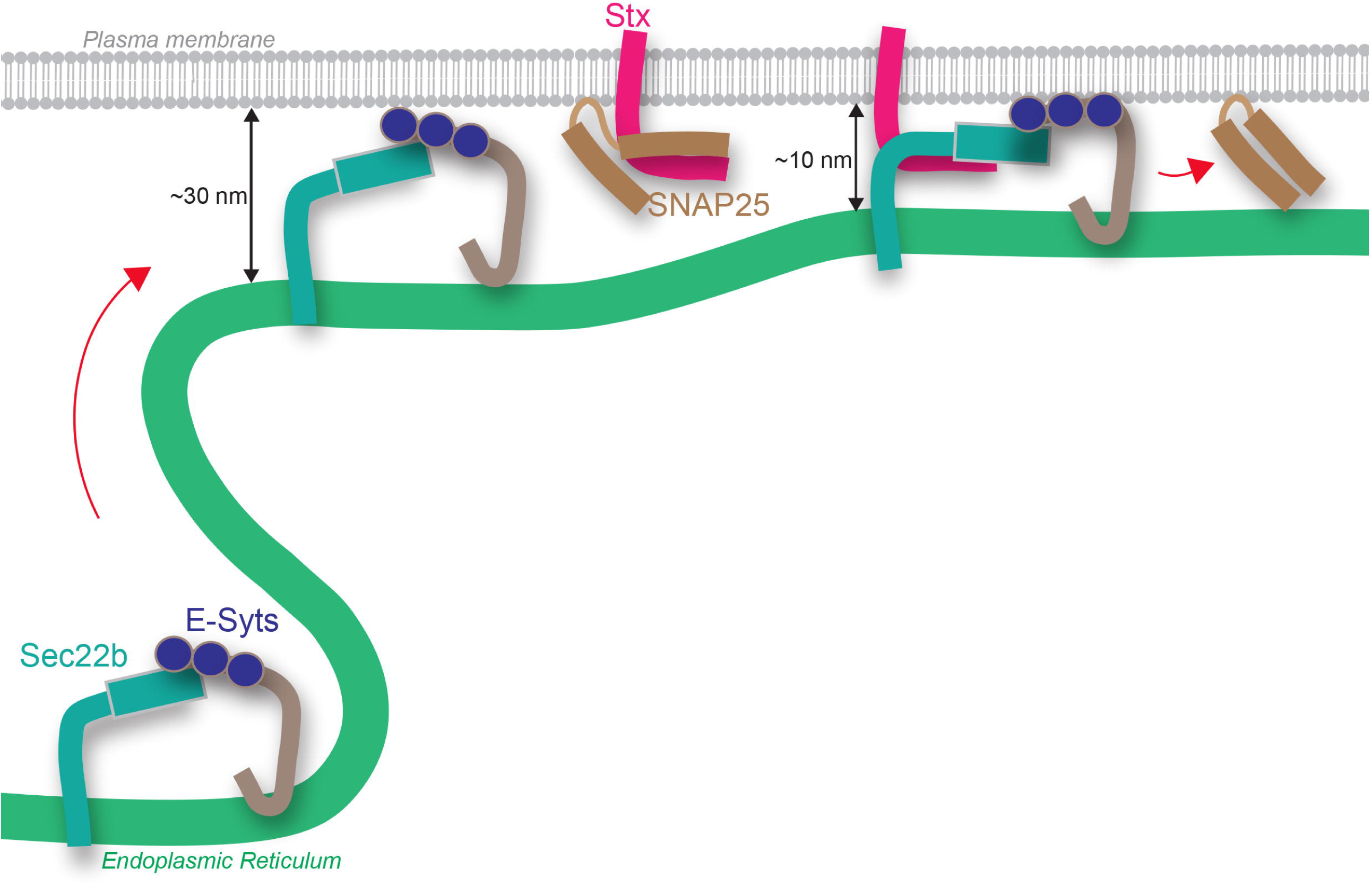
Hypothetical model for the formation of an E-Syt/SNARE-mediated ER-PM contact site. Sec22b diffuses within the ER membrane and is stabilized in the cortical ER after binding to E-Syt where it interacts with the PM-resident Stx1, generating a tight ER-PM junction (∼ 10 nm). Sec22b association to the PM-resident Stx1-SNAP-25 complex causes SNAP-25 displacement from Stx1 and potentially E-Syt, leading to the formation of a non-fusogenic assembly of Sec22b, Stx1 and E-Syts.

### Interdependence of E-Syts and Sec22b-Stx complexes for the establishment of ER-PM junctions operating in membrane expansion

The association of Sec22b to PM Syntaxins was dependent upon E-Syts expression because it was increased upon E-Syts overexpression and reduced in the absence of the three E-Syts isoforms. By promoting Sec22b/PM Stx interaction, E-Syts may increase the abundance of close contact sites between ER and PM because SNARE complexes mediate a ∼10nm distance. This shortening of the distance between the ER and the PM may further enhance the LTP activity of E-Syts as it was shown using DNA-origami in vitro (Bian *et al*, 2019). In addition, E-Syts interaction may take Sec22b away from its main function of mediating fusion events within the anterograde and retrograde membrane trafficking between ER and Golgi, in association with SNARE partners such as Stx5 or Stx18 (Burri *et al*, 2003; Liu & Barlowe, 2002). As an ER-resident protein, Sec22b can diffuse over the entire ER network, and thus is expected to visit areas of cortical ER where it can be trapped by E-Syts eventually engaged in an ER-PM tethering (Fig. 9). This in turn increases the probability to bind the PM-residing Syntaxins and form a complex in *trans*-configuration. In this view, E-Syts would have a dual function. Firstly, they would be responsible for promoting the formation of a non-fusogenic membrane-tethering complex with PM containing Sec22b and PM Stx. Secondly, the lipid transfer activity E-Syts would be enhanced by the formation of the minimal ternary complex E-Syts-Sec22b-Stx because the distance between the ER and the PM would be smaller at these interaction sites. This dual activity of E-Syts could be viewed as the generation of a MCS specialized for lipid transfer functioning in membrane growth.

### Morphogenetic effect of E-Syt overexpression

Our data showed that the elicited membrane expansion promoted by elevated E-Syt2 expression was abolished in cells expressing deleted non-functional versions of E-Syt2, i.e. E-Syt2 ΔSMP and E-Syt2 ΔMSD. The increased neurite growth and actin-rich filopodia formation induced by E-Syt2 overexpression depended on the presence of a functional Stx1 in neurons because it was prevented by treatment with BoNTC1 but not BoNTs A. This morphological phenotype also depended on Sec22b because expression of 33P mutant and the Longin domain alone both reversed the effect of E-Syt2 overexpression and BoNTD, which cleaves VAMP2, had not effect. It is thus very clear that E-Syts, Stx1 and Sec22b interact both biochemically and functionally. Interestingly, growth was not impaired in cells expressing E-Syt2 mutants, as we could not detect significant differences in membrane expansion as compared to control cells. Therefore, expression of these mutants did not act in a dominant negative manner. These results suggest that other LTPs share redundant function with E-Syts. Furthemore, *Esyt* triple knockout mice display no major defects in neuronal development and morphology (Sclip *et al*, 2016) whereas recent data showed that *dE-Syt* knockout in the fly led to major growth defect (Nath *et al*, 2019). Taken together, these evidence suggest that a functional redundancy exists among LTPs in promoting membrane growth in mammals, as the removal of one class of such proteins can be compensated by the activity of the others. The precise contribution of each class LTPs, as well as their mutual interplay, will require further studies. The effect of E-Syt overexpression suggest that LTP may be limiting factors and that fine tuning of their expression level may be critical for their function. FGFR was shown to regulate the expression of E-Syt2 in *Xenopus* embryos (Jean *et al*, 2010). Therefore, it will be particularly interesting to search if growth factors which promote axonal filopodia formation (Menna *et al*, 2009) might also control the expression level of LTPs such as E-Syts. Since actin instability is important for axon formation (Bradke & Dotti, 1999), and actin dynamics is regulated by lipidic rafts (Caroni, 2001), it will also be particularly interesting to characterize the lipidic composition of E-Syt-generated filopodia plasma membrane.

In conclusion, the protein complex between E-Syts and Sec22b unraveled here appears as an important model for further studies aimed at understanding how lipid transfer at MCS between the ER and PM could participate to the development of the neuronal cell shape. As exemplified in the case of other SNAREs like VAMP7 (Daste *et al*, 2015), our results point to a central regulatory function of the Longin domain of Sec22b.

## Materials and Methods

### Antibodies

Primary antibodies used in this study were: mouse monoclonal anti-Sec22b 29-F7 (Santa Cruz, sc-101267); rabbit polyclonal anti-ESYT2 (Sigma-Aldrich, HPA002132); rabbit polyclonal anti-Calnexin (Cell Signaling, 2433S); mouse monoclonal anti-GFP (Roche, 11814460001); mouse monoclonal anti-c-myc 9E10 (Roche, 11667203001); mouse monoclonal anti-FLAG M2 (Sigma-Aldrich); mouse monoclonal anti-Stx1 HPC-1 (Abcam, (ab3265); mouse monoclonal anti-SNAP25 (Synaptic Systems, 111011); rabbit polyclonal anti-VAMP2 (Synaptic Systems, 104 202); rabbit polyclonal anti-beta3 tubulin (Synaptic Systems, 302 302); chicken polyclonal anti-MAP2 (AbCam, ab5392); goat polyclonal anti-GFP (AbCam, ab6673); Alexa Fluor™ 633 Phalloidin (ThermoFisher, A22284). Rabbit polyclonal anti-Sec22b (TG51) and rabbit polyclonal anti-Syntaxin3 (TG0) were produced in the laboratory.

### Plasmids and cDNA clones

Plasmids encoding eGFP-ESyt2, eGFP-ESyt3, myc-ESyt2 and myc-ESyt3 were obtained from P. De Camilli (Giordano *et al*, 2013). mCherry-Sec22b, the pHluorin-tagged forms of Syntaxin1, Sec22b, and VAMP2 and GFP-tagged forms of Sec22b-P33 mutant, of Sec22b-Longin (Ribrault *et al*, 2011; Petkovic *et al*, 2014), of VAMP4 and of SNAP-25 (Mallard *et al*, 2002; Martinez-Arca *et al*, 2000) were previously described. cDNA clones of human Sec22b and Syntaxin2 and Syntaxin3 were obtained from Dharmacon (GE Healthcare) (Sec22b: Clone3051087, Accession BC001364; Syntaxin2: Clone 5296500, Accession BC047496; Syntaxin3: Clone 3010338, Accession BC007405).

### Construction of ΔLonginSec22b -pHluorin

ΔLonginSec22b-pHluorin was generated from ERS24-pQ9 construct and cloned into pEGFPC1-pHluorin using the following primers:

Forward NheI Sec22deltaLongin,

5’-CGCGCTAGCATGCAGAAGACCAAGAAACTCTACATTGAT

Reverse EcoRI Sec22deltaLongin,

5’-CGGGAATTCCAGCCACCAAAACCGCACATACA

### Construction of 3XFLAG-Secc22b, and EGFP-Syntaxin3

cDNAs of human Sec22b, and Syntaxin3 ORFs were amplified by PCR using the following primers:

Forward EcoRI-Sec22b,

5’-CAAGCTTC GAATTC ATGGTGTTGCTAACAATGATC

Reverse BamHI-Sec22b,

5’-TCCGATTCTGGTGGCTGTGAGGATCCACCGGTCG

Forward SalI-Syntaxin3,

5’-ACCG GTCGAC ATGAAGGACCGTCTGGAGCAG

Reverse SacII-Syntaxin3,

5’-GACTTTCCGTTGGGCTGAATTAACCGCGGCGGT

PCR products were ligated into the p3XFLAGCMV10 (Sigma-Aldrich) and pEGFP-C1 (Clontech) vectors, respectively to generate 3XFLAG-Sec22b, EGFP-Syntaxin2 or EGFP-Syntaxin3.

### Construction of ΔSMP myc-ESyt2 and ΔMSD myc-ESyt2

ΔSMP myc-ESyt2 and ΔMSD myc-ESyt2 mutants were generated by site-directed mutagenesis by deleting respectively fragment [119-294] and [1-72] using the following primers:

Forward DeltaSMP,

5’-CTGGGTTCATTTTCCAGACACTGAAAGTGAAGTTCAAATAGCTCAGTTGC

Reverse DeltaSMP,

5’-CCAACTGAGCTATTTGAACTTCACTTTCAGTGTCTGGAAAATGAACCCAG

Forward DeltaMSD,

5’-AGATCTCGAGCTCAAGCTTCGAATTCTCGCAGCCGCGGCCTCAAG

Reverse DeltaMSD,

5’-CTTGAGGCCGCGGCTGCGAGAATTCGAAGCTTGAGCTCGAGATCT

### Cell culture, siRNA and transfection

HeLa cells were cultured in DMEM containing GlutaMax (ThermoFisher 35050038) and supplemented with 10% (v/v) Bovine Fetal Calf Serum (Biosera 017BS346).) and 1% (v/v) penicillin/streptomycin (ThermoFisher 15140122) at 37°C and 5% CO_2_. Transfections were carried out with Lipofectamine 2000 transfection reagent (ThermoFisher 11668027) according to manufacturer’s instructions. For knockdowns, HeLa cells were transfected with control or E-Syts siRNA oligos by using Oligofectamine transfection reagent (ThermoFisher 12252011) according to manufacturer’s instructions. Double-stranded siRNAs targeting the three human E-Syts and control siRNAs were as described (Giordano *et al*, 2013). Routinely, transfected cells were cultured for 24 or 48 hours on coverslips prior to analysis.

PC12 cells were grown at 37°C and 5% CO_2_ in RPMI containing and 10% (v/v) horse serum (ThermoFisher 26050088), 5% (v/v) Bovine Fetal Calf Serum (Biosera 017BS346) and 1% (v/v) penicillin/streptomycin. Cells were coated on plastic dishes with a 1 µg/ml collagen (Sigma C7661) solution. Then, cells were differentiated with hNGF-beta (Sigma-Aldrich N1408) at 50 ng/ml for 3–4 days. PC12 transfections were carried out with Amaxa Nucleofection Kit V (Lonza VCA-1003) according to manufacturer’s instructions.

Hippocampal neurons from embryonic rats (E18) were prepared as described previously (Dotti *et al*, 1988) and modified (Danglot *et al*, 2012). Cells were grown on onto poly-L-lysine-coated 18-mm coverslips (1 mg/ml) or 30-mm plastic dishes (0.1 mg/ml) at a density of 25,000-28,000 cells/cm^2^ in Neurobasal-B27 medium previously conditioned by a confluent glial feeder layer [Neurobasal medium (ThermoFisher 21103049) containing 2% B27 supplement (ThermoFisher A3582801), and 500 μM L-Glutamine (ThermoFisher 25030024)]. Neurons were transfected before plating by using Amaxa Rat Neuron Nucleofection Kit (Lonza VPG-1003) following manufacturer’s instructions. After 3 days *in vitro* (3DIV), neurons were processed for immunofluorescence or lysed for immunoblot assays.

### Botulinum Neurotoxins (BoNTs) treatment

BoNT/A, BoNT/C,, BoNT/D were produced as previously (Strotmeier *et al*, 2011). Toxins working concentration at 1 nM in culture media were prepared from 4 µM stock solutions.

Hippocampal neurons overexpressing Myc-E-Syt2 were treated with BoNT/A, BoNT/C, BoNT/D or naïve culture media at 3DIV and maintainedfor 4 hours at 37°C and 5% CO_2_. After extensive washing with culture media, cells were processed for immunofluorescence or lysed for immunoblot assays.

### Immunofluorescence

HeLa cells and hippocampal neurons at 3DIV were fixed on coverslips in 4% paraformaldehyde in PBS for 15 minutes and quenched in 50 mM NH_4_Cl in PBS for 20 min. Cells were then permeabilized in 0.1% (w/v) Triton X-100 in PBS for 5 min and blocked in 0.25% (w/v) fish gelatin in PBS for 30 min. Primary antibodies were diluted in 0.125% fish gelatin in PBS and incubated overnight at 4°C. After washing, secondary antibodies conjµgated with Alexa 488, 568, 594 or 647 were incubated for 45 min at room temperature before mounting in Prolong medium (ThermoFisher P36930).

### Surface staining

Hippocampal neurons at 3DIV were placed on an ice-chilled metallic plate. Neurobasal medium was replaced with ice cold DMEM with 20 mM Hepes containing primary antibody (mouse anti-GFP). Cells were incubated for 5-10 min on ice. Cells where then washed with PBS at 4°C and fixed with 4% para-formaldehyde sucrose for 15 min at room temperature. Following fixation, cells were subjected to the already described whole cell staining protocol. The total pool of tagged proteins was detected with goat anti-GFP antibody. Images were acquired on the epifluorescent microscope with the same exposure in all conditions. Presence at the plasma membrane was expressed as ratio between total and surface signal.

### In situ proximity ligation assays (PLA)

In situ proximity ligation assays (PLA) to quantify protein vicinity in HeLa cells and in neurons on coverslips as indicated in figures were performed using the Duolink In Situ Detection Reagents Orange kit (Sigma-Aldrich, DUO92007). Cells were fixed, permeabilized as described above, blocked in Duolink Blocking solution (supplied with the kit) for 30 min at 37°C in a humidified chamber. This was followed by incubation with the primary antibodies rabbit Stx3 (4µg/ml), or rabbit E-Syt2 (2µg/ml) or rabbit Calnexin (2µg/ml) and mouse Sec22b (2µg/ml). For the rest of the protocol the manufacturer’s instructions were followed. Briefly, cells were washed in kit Buffer A 3 times for 15 minutes and incubated with the PLA probes Duolink In Situ PLA Probe Anti-Mouse PLUS (Sigma-Aldrich, DUO92001) and Duolink In Situ PLA Probe Anti-Rabbit MINUS (Sigma-Aldrich, DUO92005) for 1 hour at 37°C in a humid chamber followed by two washes of 5 min in Buffer A. The ligation reaction was carried out at 37°C for 30 min in a humid chamber followed by two washes of 5 min in Buffer A. Cells were then incubated with the amplification-polymerase solution for 100 min at 37°C in a dark humidified chamber. After two washings with kit Buffer B for 10 minutes followed by a 1 minute wash with 0.01x Buffer B, cells were mounted using the Duolink In Situ Mounting Medium with DAPI (Sigma-Aldrich, DUO92040).

### Microscopy and image analysis (missing STED)

#### Confocal imaging

Z-stacked confocal images of neurons and HeLa cells were acquired in a Leica TCS SP8 confocal microscope (Leica Microsystems CMS GmbH), using a 63x/1.4 Oil immersion objective.

#### HeLa cells live imaging

HeLa co-expressing mCherry-Sec22b and GFP-E-Syt2 were transferred to an imaging chamber (Chamlide EC) and maintained in Krebs-Ringer buffer (140mM NaCl, 2.8mM KCl, 1mM MgCl2, 2mM CaCl2, 20mM HEPES, 11.1mM glucose, pH 7.4). Time-lapse videos were recorded at 5s intervals for 2min using an inverted DMI6000B microscope (Leica Microsystems) equipped with a 63X/1.4-0.6 NA Plan-Apochromat oil immersion objective, an EMCCD digital camera (ImageEMX2, Hamamatsu) and controlled by Metamorph software (Roper Scientific, Trenton, NJ). To virtually abrogate latency between two channels acquisition, illumination was sequentially provided by a 488nm- and a 561nm-diode acousto-optically shuttered lasers (iLas system; Roper Scientific) and a dualband filter cube optimized for 488/561nm laser sources (BrightLine; Semrock) was used. Environmental temperatures during experimental acquisitions averaged 37°C. FIJI software was used for bleaching correction and for movies montage. Binary mask of particles was generated by applying the wavelet-based spot detector plugin of Icy imaging software (http://icy.bioimageanalysis.org) to each channel sequence.

#### STED imaging

Neuronal membrane was labeled on live neurons using the lectin Wheat Germ Agglutinin (WGA) coupled to Alexa 488 nm for 10 min at 37°C. Neurons were washed and fixed with a 4% PFA /0,2% Glutaraldehyde mixture and then processed for immunochemistry. Growth cone were imaged with 3D STED microscopy using the 775 nm pulsed depletion laser. Depletions were carried out on primary antibody to endogenous E-Syt2 labelled with secondary antibodies coupled to Alexa 594. Sec22b-pHL were labelled with primary anti-GFP and ATTO647N secondary antibodies. Acquisitions were done in 3D STED so that the voxel size were isotropic. Typically, we imaged a matrix of 1400×1400 pixels over 20 to 25 z planes to include all the growth cone volume. Growth cone was segmented using Icy “HK means” plugin (http://icy.bioimageanalysis.org/plugin/hk-means/). Spatial distribution analysis of E-Syt2-and Sec22 was done using “Icy SODA” plugin (Lagache *et al*, 2018) and a dedicated LD protocol automation. Briefly, E-Syt2 and Sec22b distribution were analysed through the Ripley function. Statistical coupling between the two molecules was assessed in concentric target. Over 13784 E-Syt2 clusters analysed in 4 different growth cones, 20 percent were statistically associated with Sec22b. When associated, the 2 molecules were at 84nm ±9nm, which corresponds to 34% of Secc22b clusters. This coupling is of very high significance since the p value was ranging between 10-5 and 10-24. Distance to the plasma membrane (d) was measured using the Icy ROI inclusion analysis plugin (Lagache *et al*, 2018).

#### PLA signal

Maximum intensity projections of a confocal z-stack including a whole cell were performed to observe the maximum amount of PLA puncta. The number of puncta per cell was counted using the Cell Counter plugin in Fiji/ImageJ. In neurons, the PLA puncta were separately counted in cell body, neurites and growth cones and the PLA signals were divided by the area of the coresponding compartments.

#### Neurite length

For the analysis of neurite length of cultured neurons, images were analyzed using the NeuronJ plugin in Fiji/ImageJ on the maximal intensity projections of z-stacks of the eGFP channel. Main process and branches were measured for each neuron. We could not detect any association among individual cells thereby we considered each cell a sampling unit.

#### Area of spikes

in Myc-E-Syt2 overexpressing HeLa cells, the spikes area was measured on the maximal intensity projections of z-stacks of the eGFP channel. The measurement was carried by subtracting the Region on Interest (ROI) of the cell without spikes, obtained with the tool Filters/Median on Fiji/ImageJ by applying a radius of ∼80 pixels, from the ROI of the entire cell.

### GFP-Trap pull-down and Immunoblotting

Transfected HeLa or PC12 cells were lysed in TBS (20mM Tris-HCl, pH7.5, 150mM NaCl), containing 2mM EDTA, 1% Triton-X100 and protease inhibitors (Roche Diagnostics). of clarified lysate was obtained by centrifugation at 16,000xg 15 min, and 1 mg protein was submitted to GFP-Trap pull-down for 1 hour at 4 °C under head-to-head agitation using 10μl of Sepharose-coupled GFP-binding protein (Rothbauer *et al*, 2008) prepared in the lab After four washes with lysis buffer, beads were heated at 95°C for 5 min in reducing Laemmli sample buffer. Soluble material was processed for SDS-PAGE using 10 % acrylamide gels and transferred on nitrocellulose membranes (GE-healthcare). The membranes were blocked with 2.5% (w/v) skimmed milk, 0.1% (w/v) Tween-20 in PBS. Membrane areas of interest were incubated with primary antibodies as indicated in figure legends. After washing, the membranes were blotted with HRP-coupled secondary antibodies. Revelation was carried out by using a ChemiDoc luminescence imager (BioRad).

### Statistical analysis

Calculations were performed in Microsoft Excel. GraphPad Prism software were used for statistical analyses. For each dataset, at least three independent experiments were considered and all data are shown as mean ± SEM. Data were analyzed using one-way ANOVA followed by Dunn’s, Tukey or Dunnet post-hoc tests were applied as indicated in figure legends.

## Acknowlegements

We thank all members of the Galli laboratory for their assistance and discussions. Work in our group was funded by grants from Association Française contre les Myopathies (Research Grant 16612), the French National Research Agency (*NeuroImmunoSynapse* ANR-13-BSV2-0018-02; *MetDePaDi* ANR-16-CE16-0012), the Institut National du Cancer (PLBIO 2018-149), the Fondation pour la Recherche Médicale (FRM), *Who am I?* Labex (Idex ANR-11-IDEX-0005-01), awards of the Fondation Bettencourt-Schueller to T.G. We acknowledge the IPNP Neurimag Imaging facility and are deeply grateful to Leducq foundation for the grant to TG supporting the acquisition of the NeurImag SP8 3DSTED confocal system.

## Figure Legends

**Figure S1. The Sec22b**Δ**Longin mutant reaches the cell surface**

(A-B) Hippocampal neurons at 3DIV were transfected with Sec22b or Sec22bΔL mutant tagged with pHluorin at the C-terminal, so that if the Sec22b compartment fuses with the PM, it would be detected by anti-GFP antibody in the medium. N-terminally tagged Sec22b (GFP-Sec22b) has been used as a negative control since its GFP tag should never be exposed extracellularly and detected with anti-GFP antibody, even in the event of membrane fusion. VAMP2-pHL was used as positive control for fusion. For surface staining anti-GFP antibody was allowed to bind for 10 min at 4°C, neurons were fixed and processed for immunochemistry. (A) Scheme of the topology of constructs in the PM if fusion is assumed to occur. (B, left) Representative images. Scale bar, 10 µm (B, right) Quantitative analysis of surface staining versus total staining expressed as normalized ratio to GFP-Sec22b. Oneway ANOVA P<0,0001 ***, Dunn’s multiple comparison post-test labeled on graph. (C) Representative images of COS7 cells expressing Sec22b-HA (left) or Sec22bΔL-HA (right) and labeled with an anti-HA antibody. Note that the typical reticular localization of Sec22b is partially lost for the Sec22bΔL mutant.

**Figure S2. Actin-populated filopodia display clusters of E-Syt2**

Representative images of a 3DIV hippocampal neuron labeled for Myc-E-Syt2 (red) and actin (phalloidin labeling, green). Actin is present in filopodia resulting by Myc-E-Syt2 overexpression. In the inset, the immunostaining of Myc-E-Syt2 clearly reveals puncta (arrowheads) which are present in thin actin-positive filopodia.

**Movie 1. Dynamics of mCherry-Sec22b and GFP-E-Syt2 in HeLa**

Live-microscopy recording of mCherry-Sec22b and GFP-E-Syt2 movements in HeLa cells. The inset shows a region where the peripheral colocalized molecules have a notably reduced motility as compared to material with less peripheral distribution.

