## Supplementary figures and images for "The interaction between non-fusogenic Sec22b-Stx complexes and Extended-Synaptotagmins promotes neurite growth and ramification"

### Supp Figure 1

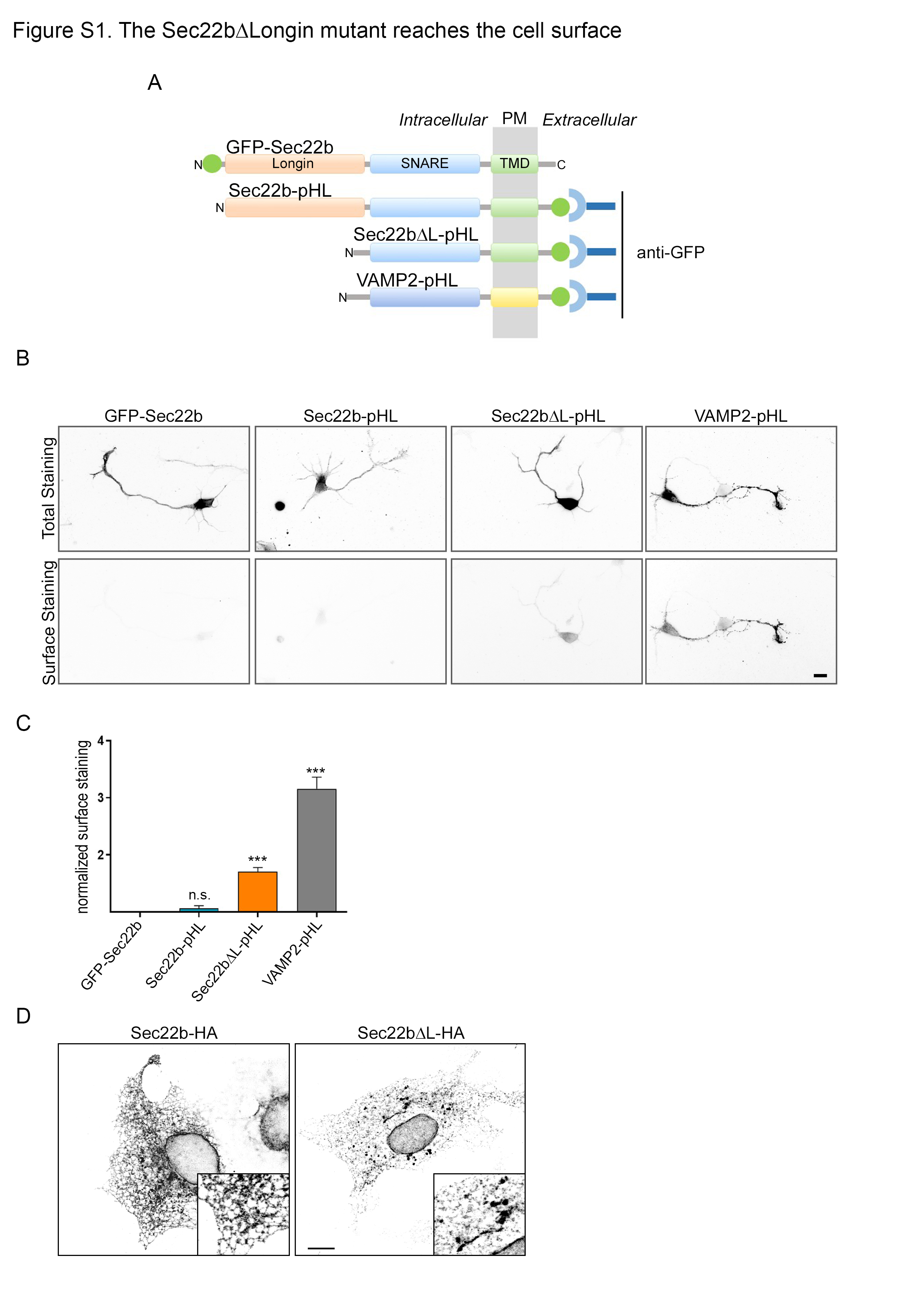

### Supp Figure 2

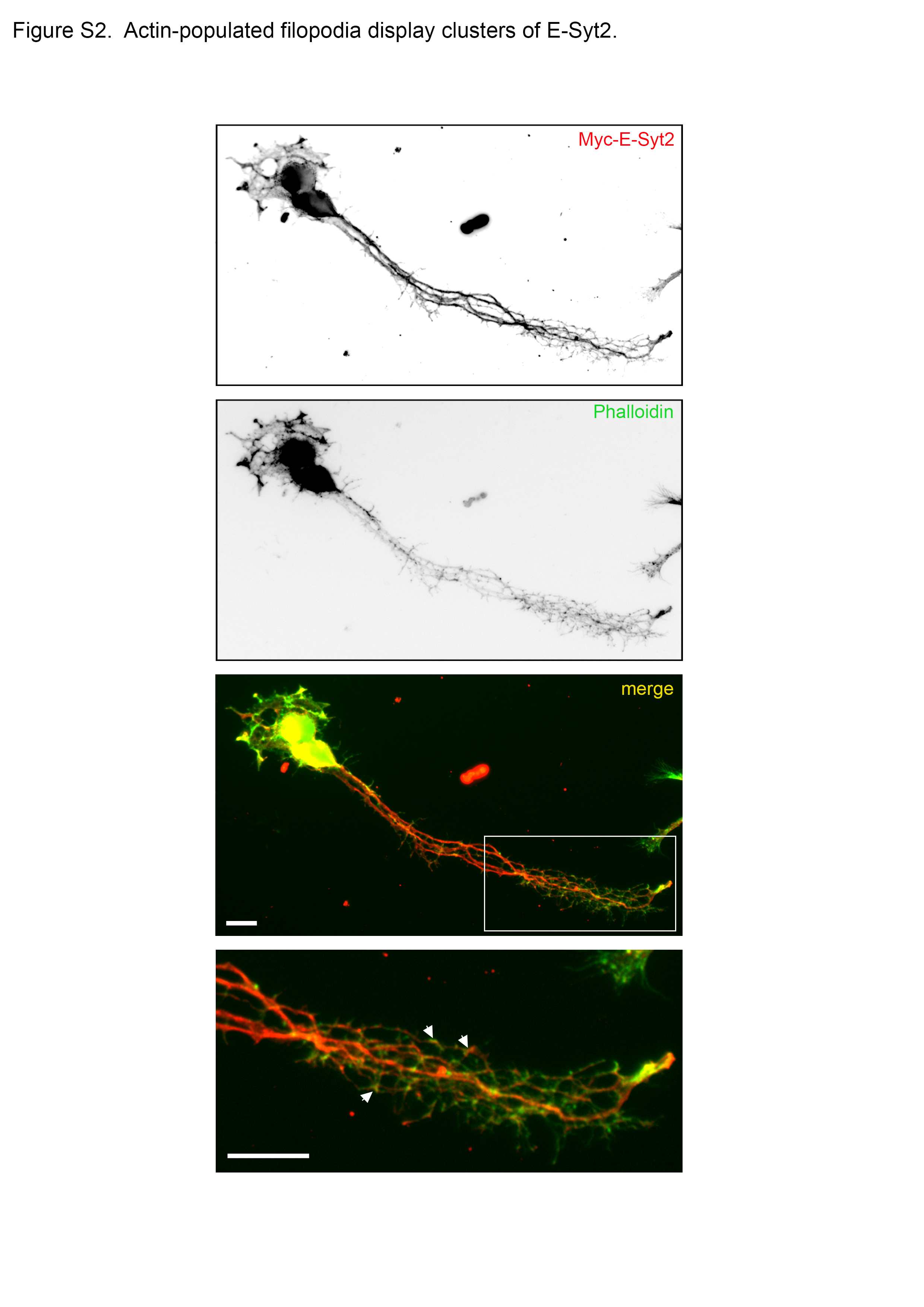
